# Notch signaling in tumor vasculature programs cancer-associated fibroblasts to suppress anti-tumor immunity

**DOI:** 10.1101/2022.02.16.480755

**Authors:** Yu Zhu, Menglan Xiang, Kevin F. Brulois, Nicole H. Lazarus, Junliang Pan, Eugene C. Butcher

**Affiliations:** Laboratory of Immunology and Vascular Biology, Department of Pathology, Stanford University School of Medicine, Stanford, CA, USA; Palo Alto Veterans Institute for Research, Veterans Affairs Palo Alto Health Care System, Palo Alto, CA, USA

## Abstract

Scarcity of tumor-infiltrating T cells poses significant challenges to cancer treatment, but mechanisms that regulate T cell recruitment into the tumor microenvironment are unclear. Here we ask if the endothelial lining of tumor vasculature suppresses T cell infiltration. Using mouse pancreatic ductal adenocarcinoma models, we found that Notch signaling in endothelial cells (ECs) inhibits the pro-inflammatory functions of cancer-associated fibroblasts (CAFs) and prevents CAFs from secreting CXCL10, a chemokine that recruits anti-tumor T cells via its receptor CXCR3. Abrogation of canonical Notch signaling in ECs reprogrammed the phenotype of CAFs from myofibroblasts into pro-inflammatory fibroblasts, unleashed interferon gamma (IFNγ) responses in the tumor, and stimulated CXCL10/CXCR3-mediated recruitment of T cells to inhibit tumor growth. Collectively, these data uncover an important role of endothelial Notch signaling in shaping the tumor immune microenvironment, and suggest the potential of targeting EC-CAF crosstalk as an approach to enhance anti-tumor immunity in immunologically cold tumors.

**In brief:** How blood vasculature shapes the tumor immune microenvironment is poorly defined. This study demonstrates that tumor endothelial cells reprogram cancer-associated fibroblasts to limit anti-tumor T cell recruitment, and suggests the potential of targeting endothelium-fibroblast crosstalk to overcome T cell scarcity in “cold” tumors and enhance anti-tumor immunity.

## Introduction

Immune cells within the tumor microenvironment play a key role in the regulation of cancer development (Binnewies et al., 2018; Fridman et al., 2012; Galon et al., 2006; Hanahan and Coussens, 2012; Zhang et al., 2003). T cells in particular display anti-tumor activities that could be manipulated for cancer therapy. In animal models and cancer patients, CD8+ cytotoxic T cells are essential for tumor inhibition by inducing malignant cell apoptosis in many types of cancers. Th1 polarized CD4+ T-helper cells are also important in orchestrating anti-tumor responses (Borst et al., 2018). However, many tumors fail to recruit tumor-infiltrating CD8+ and Th1 CD4+ T cells (Joyce and Fearon, 2015; Thommen and Schumacher, 2018; van der Woude et al., 2017), resulting in an immunologically “cold” microenvironment that shields tumor cells from immune surveillance, thus allowing tumors to survive, proliferate, and metastasize. This poses a significant challenge to cancer treatment. However, mechanisms underlying the scarcity of anti-tumor T cells are not well understood.

Recruitment of T cells to the tumor tissue is a tightly regulated process dependent on blood vasculature (Butcher and Picker, 1996; Del Prete et al., 2017; Nourshargh and Alon, 2014). Endothelial cells (ECs) lining the blood vessels are essential in mediating T cell trafficking under physiological and pathological conditions (Zabel et al., 2015). Our laboratory and others have uncovered molecular programs by which blood ECs bind circulating leukocytes and mediate immune cell trafficking (Butcher, 1991; Kunkel and Butcher, 2002; Springer, 1994). However, it is unclear whether tumor ECs interact with other stromal components of tumor tissues to limit T cell infiltration and shape the local immune milieu.

In this study, we interrogated the role of endothelial Notch signaling in shaping the tumor microenvironment. Notch signaling regulates many activities and functions of the tumor vasculature, including angiogenic sprouting, proliferation, and arterial-venous specification (Adams, 2003; Hellstrom et al., 2007; Ley et al., 2007; Mack and Iruela-Arispe, 2018; Ntziachristos et al., 2014; Phng and Gerhardt, 2009; Rehman and Wang, 2006). Therapies blocking endothelial Notch have been shown to promote defective angiogenesis and inhibit tumor growth (Noguera-Troise et al., 2006; Ridgway et al., 2006). However, it is not well established whether Notch signaling in ECs sculpts the tumor immune landscape. Unexpectedly, we uncovered that tumor-associated ECs interact with cancer-associated fibroblasts (CAFs) to inhibit the recruitment of anti-tumor T cells by suppressing CAF production of CXCL10, a chemokine necessary for CXCR3-mediated recruitment of CD8+ and Th1 CD4+ T cells. Endothelial Notch signaling is essential for this crosstalk. EC-specific abrogation of the master transcription factor of canonical Notch signaling, Rbpj, reprogrammed the phenotype of CAFs from myofibroblasts into pro-inflammatory CAFs that better support immune responses. Abrogation of Notch in tumor ECs also unleashed the production of interferon gamma in the tumor microenvironment, which enhanced CXCL10 production by CAFs to promote T cell homing in a CXCR3-dependent manner to ultimately inhibit tumor growth.

## Results

### Notch signaling is upregulated in blood vessels to promote PDAC growth

Human pancreatic ductal adenocarcinoma (PDAC) often display limited T cell infiltration, which is recapitulated by many mouse PDAC models (Hingorani et al., 2005; Jiang et al., 2016). To determine how blood vessels regulate immune cell recruitment and shape the tumor environment, we established orthotopic PDAC tumors using the KP1 cell line derived from autochthonous tumors from p48-Cre, Kras^G12D^, p53^f/f^ (KPPC) mice (Jiang et al., 2016). First, we asked whether Notch signaling is active in tumor blood endothelial cells (BECs). Towards this end, we isolated blood endothelial cells (lineage-CD31+podoplanin-) cells from normal mouse pancreas and orthotopic KP1 tumors by fluorescence-activated cell sorting, and performed quantitative polymerase chain reaction (QPCR) analyses to quantify the expression of Notch target genes. Compared to the BECs isolated from normal mouse pancreas, the BECs in orthotopic tumors exhibited increased expression of *Hey1*, *Hes1*, and *Rbpj* (Figure 1A), transcriptional targets activated upon engagement of canonical Notch pathways (Kopan and Ilagan, 2009). These data suggest that BECs in the pancreas upregulate Notch signaling as the tumors develop.

**Figure 1.**
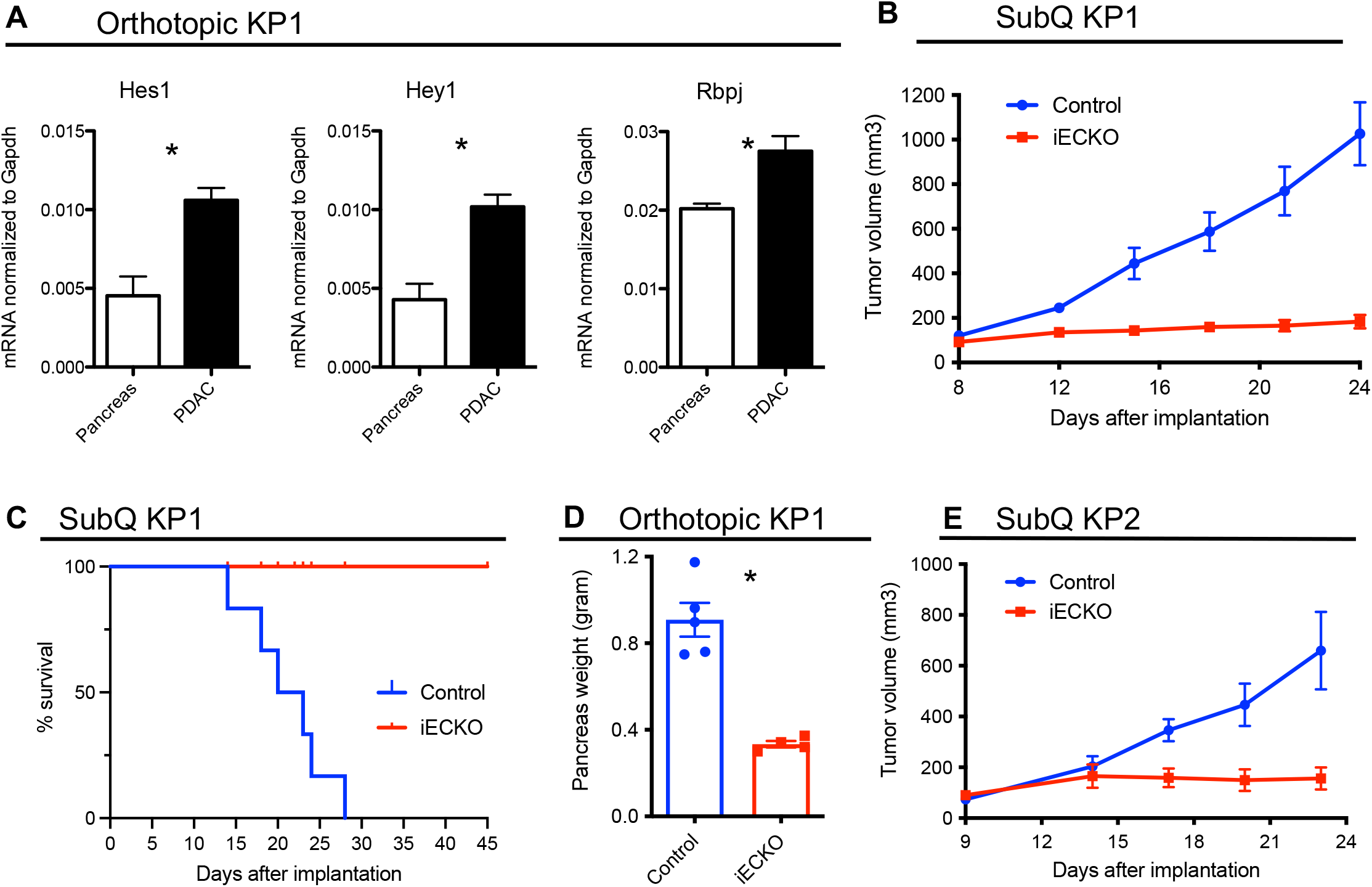
PDAC upregulates canonical Notch signaling in blood endothelial cells to promote tumor growth. A. QPCR quantification of Notch transcriptional target genes in blood endothelial cells from normal pancreas and orthotopic KP1 tumors. B. Volume of subcutaneous KP1 tumors in control and Rbpj^iECKO^ mice. C. Survival of mice bearing subcutaneous KP1 tumors. D. Weight of tumor-bearing pancreas in orthotopic KP1 tumor-bearing control and Rbpj^iECKO^ mice. E. Volume of subcutaneous KP2 tumors in control and Rbpj^iECKO^ mice.

To ask whether Notch signaling in endothelial cells was necessary for tumor growth, we genetically disrupted *Rbpj* in endothelial cells (Wang et al., 2010) using the Cdh5(PAC)-Cre^ERT2^ x Rbpj^f/f^ mice (referred to as Rbpj^iECKO^ hereafter). *Rbpj* encodes an essential transcription factor that mediates canonical Notch signaling. We established subcutaneous tumors using the KP1 cell line, and treated Rbpj^iECKO^ mice with tamoxifen to delete Rbpj when tumors reached approximately 0.5cm in length. While tumors in tamoxifen-treated control mice continued expanding in size, tumors in Rbpj^iECKO^ mice ceased growing shortly after the induction of Cre recombination (Figure 1B), at least doubling mouse survival duration (Figure 1C). Loss of Rbpj in ECs also significantly inhibited tumor growth in orthotopically implanted KP1 tumors (Figure 1D). Similar results were observed in tumors KP2, an independent PDAC cell line derived from p48-Cre, Kras^G12D^, p53^m/wt^ (KPC) mice (Figure 1E and Supplemental Figure 1). Collectively, these data suggest that canonical Notch signaling in tumor endothelial cells is necessary for tumor expansion.

### Abrogation of endothelial Notch signaling promotes the infiltration of anti-tumor T cells, but not myeloid cells

To determine whether canonical Notch signaling in endothelial cells regulates T cell access for immune responses in the tumor tissue, we deleted Rbpj in ECs of PDAC-tumor bearing mice, and profiled tumor-infiltrating leukocytes by flow cytometry. We found that Rbpj^iECKO^ did not significantly alter the abundance of tumor-associated macrophages, monocytes/monocytic myeloid derived suppressor cells (Mo-MDSCs), neutrophils/granulocytic MDSCs (G-MDSCs), or eosinophils (Figure 2A, Supplemental Figure 2). However, loss of endothelial Notch signaling significantly increased the infiltration of T cells, including CD8+ and CD25-CD4+ effector T cells, and to a lesser extent regulatory T cells (T_re__g_s) (Figure 2B). The increase in T cell infiltration occurred rapidly within 24 hours after tamoxifen induction (Supplemental Figure 3A) and was maintained throughout tumor progression. We confirmed the flow cytometric data by immunofluorescence imaging (Figure 2C). Interestingly, in PDAC tissues of Rbpj^iECKO^ mice, we observed clusters of CD8+ T cells surrounding cytokeratin-positive cells that expressed cleaved caspase 3, indicative of apoptotic tumor cells (Figure 2C). A similar increase in T cells but not of myeloid cells was also observed in orthotopic KP2 tumors (Supplemental Figures 3B).

**Figure 2.**
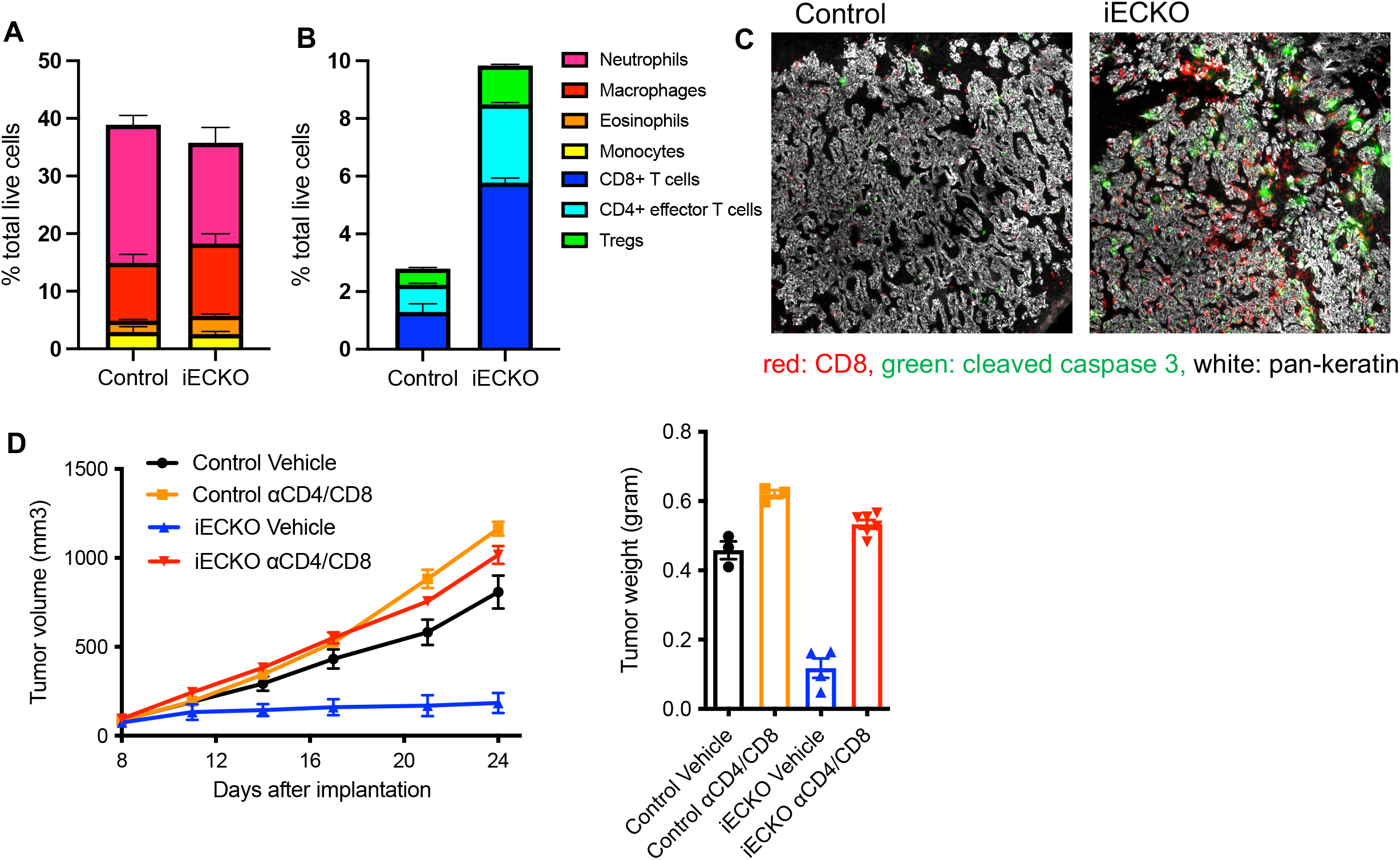
Abrogation of endothelial Notch signaling promotes the infiltration of T cells in PDAC. A-B. Flow cytometric quantification of myeloid cells (A) and T cells (B) in subcutaneous KP1 tumors of control and Rbpj^iECKO^ mice. C. Immunofluorescence imaging of CD8 and cleaved caspase 3 in subcutaneous KP1 tumors. D. Volume of subcutaneous KP1 tumors in control and Rbpj^iECKO^ mice treated with CD4/CD8 depleting antibodies.

**Figure 3.**
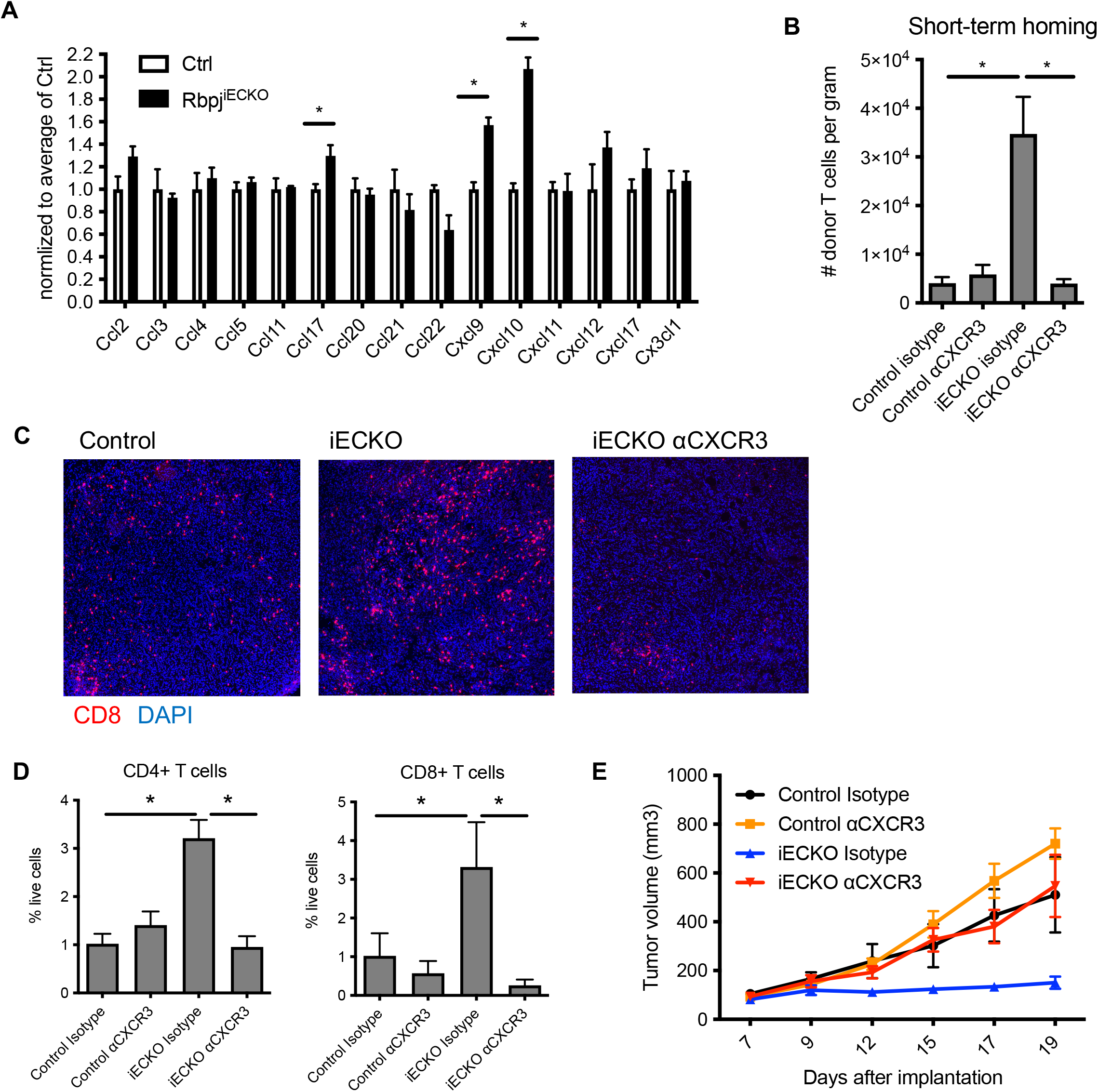
Abrogation of endothelial Notch signaling enhances CXCL10/CXCR3-mediated recruitment of anti-tumor T cells. A. QPCR quantification of chemokine array on whole tissues of subcutaneous KP1 tumors in control and Rbpj^iECKO^ mice. B. Quantification of tdTomato+ donor-derived T cells recruited to orthotopic KP1 tumors in control and Rbpj^iECKO^ recipient mice treated with anti-CXCR3 or isotype control. C. Immunofluorescence imaging of CD8+ T cells in KP1 tumors in control and Rbpj^iECKO^ mice. D. Flow cytometric quantification of T cells on tissues in (C). E. Tumor burden of mice in (C-D).

To determine whether the increased T cell infiltration was necessary for the inhibition of tumor growth seen in mice deficient in endothelial Notch, we treated tumor-bearing Rbpj^iECKO^ and control mice with CD4 and CD8 depleting antibodies. Interestingly, while loss of Rbpj in ECs halted PDAC growth, depletion of T cells unleashed tumor progression to a level comparable to that seen in the control mice (Figure 2D). T cell depletion also abolished the Rbpj^iECKO^-mediated tumor inhibition in orthotopic tumors (Supplemental Figure 3C). These data suggest that the enhanced T cell infiltration is necessary for tumor inhibition in Rbpj^iECKO^ mice.

Taken together, these findings suggest that the enhanced Notch signaling in tumor endothelial cells inhibits the infiltration of anti-tumor T cells, thereby shielding tumor cells from T cell-mediated killing. Abrogation of Notch signaling in ECs alleviated this inhibition and selectively enhanced T cell infiltration to halt cancer expansion.

### Loss of Notch signaling in ECs enhances CXCL10/CXCR3-mediated recruitment of anti-tumor T cells to PDAC

Given the observations that the infiltration of T cells, but not the myeloid cells, was selectively enhanced in Rbpj^iECKO^ tumors, we hypothesized that loss of endothelial Notch signaling led to increased production of chemokines that promote T cell recruitment. We isolated mRNA from KP1 tumor tissues established in Rbpj^iECKO^ and control mice, and performed QPCR analyses to profile the expression of an array of chemokines. We did not see an increased expression of most chemokines that we examined, including *Ccl2*, *Ccl3*, *Ccl5*, *Ccl11*, and *Cx3cl1* (Figure 3A). These are consistent with the observations that Notch deficiency in ECs did not alter the infiltration of monocytes or eosinophils. On the other hand, the expression of *Cxcl9* and, more prominently, *Cxcl10* significantly increased in the tumors of Rbpj^iECKO^ mice (Figure 3A). CXCL9 and CXCL10 mediate T cell recruitment by binding the receptor CXCR3, which is predominantly expressed on activated CD8+ T cells and Th1 differentiated CD4+ T cells (Groom and Luster, 2011b; Mach et al., 1999; Woods et al., 2017).

To determine whether CXCR3 signaling was required for T cell recruitment, we assessed short-term lymphocyte homing (Figure 3B). We established orthotopic KP2 tumors in Rbpj^iECKO^ and control recipient mice, and treated the mice with tamoxifen when the tumors became palpable. For donors we established orthotopic KP2 tumors in Rosa26-mTmG mice, whose cells carry tdTomato fluorescence. We then isolated lymphocytes from the tumor-draining lymph nodes (LNs) of the donor mice and injected them intravenously into the recipients. Concomitantly with cell transfer, mice were treated intraperitoneally with the CXCR3 blocking antibody or the isotype controls. Sixteen hours after injection, we dissociated tumors from recipients and performed flow cytometry to quantify the number of tdTomato+ donor-derived cells that homed to the tumor tissue. Consistent with our hypothesis, more donor-derived T cells homed to the tumors of Rbpj^iECKO^ mice compared to control mice; and the increased T cell homing was abolished upon CXCR3 blockade. These data suggest that CXCR3 mediates the enhanced recruitment of T cells into PDAC tumors in Rbpj^iECKO^ mice (Figure 3B).

To determine whether the CXCL9/10-CXCR3 pathway was required for Rbpj^iECKO^-mediated tumor inhibition, we treated KP1 tumor-bearing mice with CXCR3 blocking antibody or isotype control (Figures 3C–3E). Anti-CXCR3 treatment successfully blocked the induced infiltration of CD8+ and CD4+ T cells into the tumor tissue, as determined by immunofluorescence imaging and flow cytometry (Figures 3C and 3D). Importantly, CXCR3 blockade abolished the inhibition of tumor growth seen in the isotype antibody-treated Rbpj^iECKO^ mice (Figure 3E), suggesting that the tumor inhibition induced by Notch deficiency in ECs depends on CXCR3-mediated T cell infiltration. These observations were directly due to the blockade of CXCR3 on T cells, as CXCR3 was expressed predominantly on T cells, but not on tumor cells or other stromal populations (Supplemental Figure 4). Taken together, these data suggest that endothelial Notch signaling blocks the infiltration of anti-tumor T cells by suppressing the CXCL9/10-CXCR3-mediated T cell homing, thereby promoting an immunologically cold tumor microenvironment that shields tumor cells from immune attack.

**Figure 4.**
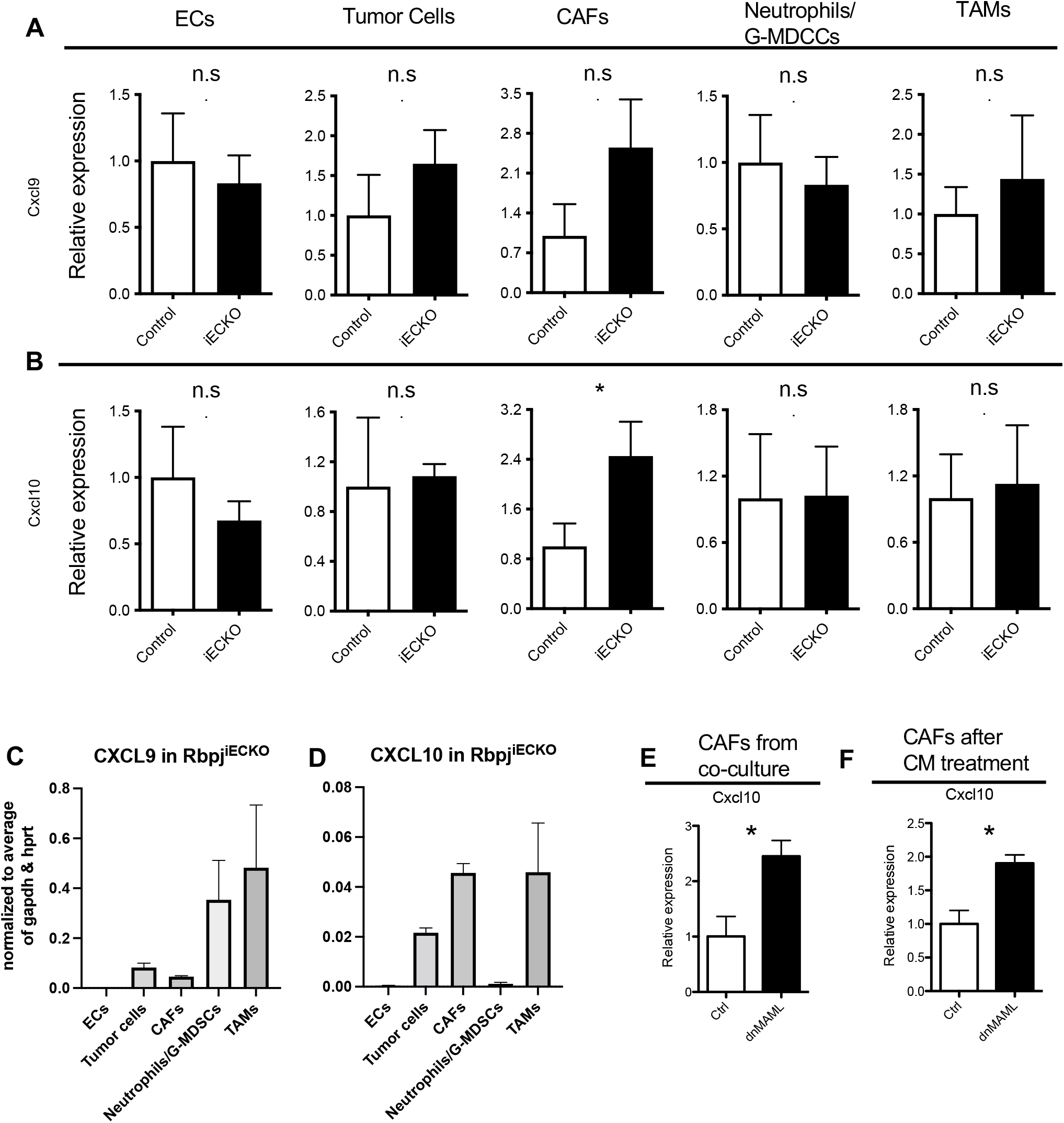
Abrogation of endothelial Notch signaling enhances CXCL10 production by cancer-associated fibroblasts. A-B. QPCR quantification of *Cxcl9* (A) and *Cxcl10* (B) transcripts in ECs, tumor cells, CAFs, neutrophils/G-MDSCs, and TAMs sorted from KP1 tumors of Rbpj^iECKO^ mice. C-D. QPCR quantification of *Cxcl9* (C) and *Cxcl10* (D) normalized against the average of Gapdh and Hprt in indicated populations of KP1 tumors from control and Rbpj^iECKO^ mice. E. QPCR quantification of *Cxcl10* transcript in CAFs co-cultured with control bEND3 or bEND3-dnMAML cells. F. QPCR quantification of *Cxcl10* transcript in CAFs treated with conditioned media from control bEND3 or bEND3-dnMAML cells.

### Loss of Notch signaling in ECs enhances the production of CXCL10 by Cancer-Associated Fibroblasts

Next, we sought to identify the sources of increased CXCL9/10 production in the tumor microenvironment. We initially hypothesized that Notch deficiency in ECs led to increased CXCL9 and CXCL10 production in a cell autonomous manner. However, when we performed QPCR analyses on sorted BECs, we found that Rbpj deficiency did not significantly change the expression of genes encoding these chemokines (Figures 4A and 4B). This suggests that the increased production of chemokines came from other cells in the tumor tissue.

To search for other cellular sources, we performed transcriptomic profiling of cells from the tumor environment of Rbpj^iECKO^ and control mice. To ensure sufficient representation of relevant cell types, we sorted BECs, tumor cells, T cells, cancer-associated fibroblasts (CAFs), natural killer (NK) cells, pericytes, and myeloid cells including TAMs, monocytes/Mo-MDSCs, neutrophils/G-MDSCs, and eosinophils (Figures 5A–5B and Supplemental Figures 2 and 5), and pooled the cells for single cell RNA sequencing (scRNAseq) analysis. *Cxcl9* and *Cxcl10* were not changed in most cell types examined, including TAMs, G-MDSCs, and tumor cells. However, in CAFs isolated from Rbpj^iECKO^ mice, we saw a significant increase of *Cxcl10* compared to CAFs from control tumor-bearing mice (Figure 5H). QPCR analyses on sorted CAFs confirmed that Rbpj^iECKO^ mice had significantly increased *Cxcl10* transcripts, and a trend of increase in *Cxcl9* (Figures 4A–4B). Of note, of all the cell types examined, CAFs and TAMs expressed the highest levels of *Cxcl10* in Rbpj^iECKO^ mice (Figure 4D). On the other hand, the predominant sources of CXCL9 are likely TAMs and G-MDSCs, but their *Cxcl9* expression was not affected by altered endothelial Notch signaling, and thus is unlikely to be responsible for the inhibited tumor growth (Figures 4A–4D). Neither chemokine was highly expressed in blood endothelial cells (Figures 4C and 4D). Taken together, these data suggest that Notch signaling in ECs suppressed fibroblast expression of CXCL10, and that abrogation of endothelial Notch signaling alleviated the repression of fibroblast CXCL10 to promote T cell recruitment.

**Figure 5.**
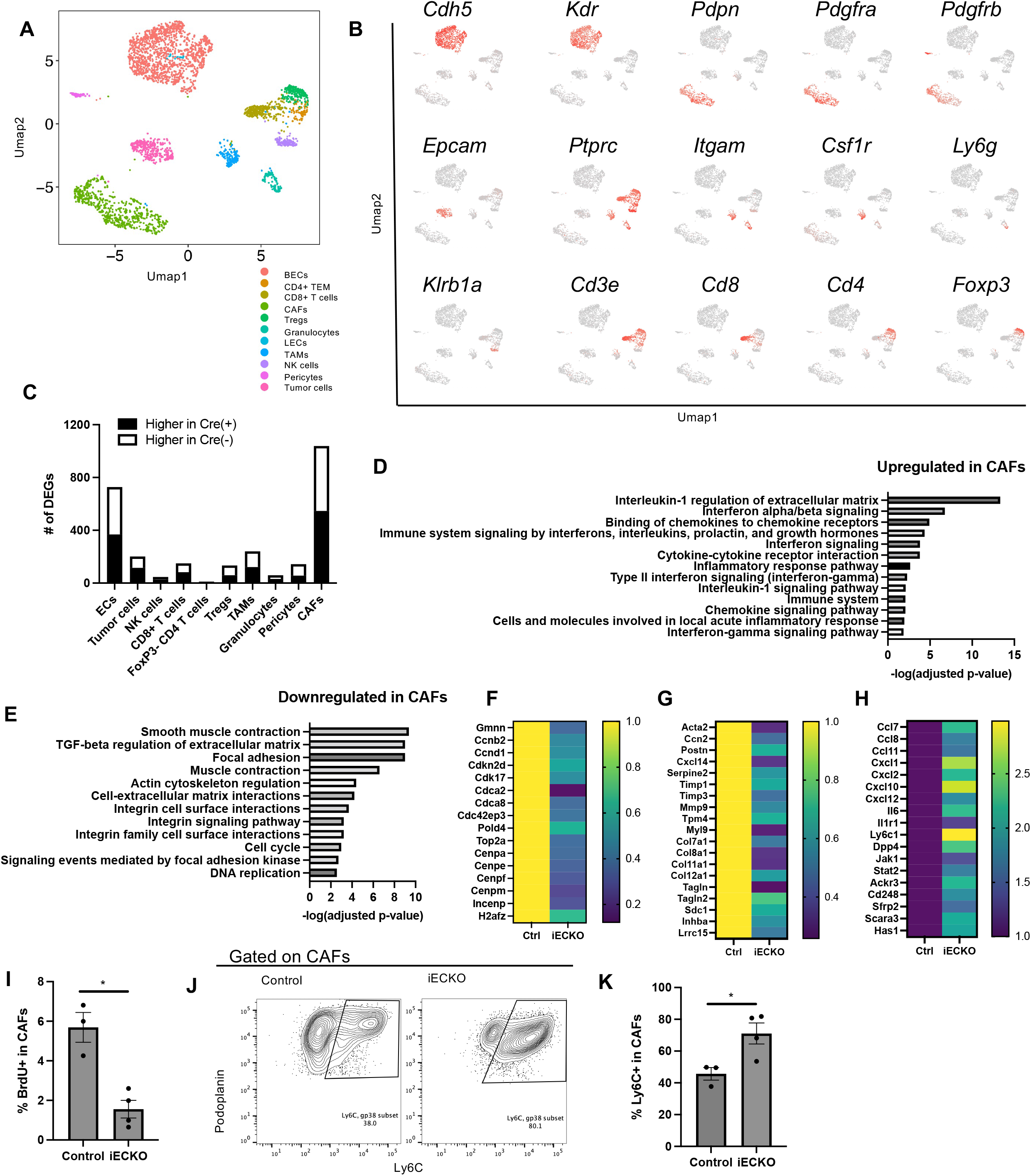
Abrogation of endothelial Notch reprograms myofibroblastic CAFs into pro-inflammatory CAFs. A. Umap plots of tumor and stromal cells processed by scRNAseq analyses. B. Expression of selected genes for cell identification. C. Number of differentially expressed genes (DEGs) in indicated cell types in response to Rbpj_iECKO_. D-E. BioPlanet pathway analyses of DEGs in CAFs that are upregulated (D) or downregulated (E) by Rbpj^iECKO^. F-I. Select CAF DEGs involved in cell cycle (F), myofibroblastic functions (G), and inflammation (H). I. BrdU analyses of CAFs from KP1 tumors in control and Rbpj^iECKO^ mice. J. Representative plots of Ly6C expression in CAFs from KP1 tumors in control and Rbpj^iECKO^ mice. K. Quantification of Ly6C+ CAFs from KP1 tumors in control and Rbpj^iECKO^ mice.

Next, we assessed whether Notch signaling in ECs could suppress the expression of CXCL10 by CAFs in an *ex vivo* system without compounding factors from the complex tumor microenvironment. Towards this end, we sorted CAFs from orthotopic KP1 tumors, and co-cultured CAFs with BECs that have intact or deficient Notch signaling. We used bEND3 cells, a microvascular cell line widely adopted for *in vitro* EC culture studies (Montesano et al., 1990). To inhibit Notch signaling, we transduced bEND3 cells with a lentivirus carrying a construct that expresses dominant negative Mastermind (dnMAML). dnMAML is a well validated and specific inhibitor of canonical Notch signaling, phenocopying Rbpj deficiency (Chiang et al., 2016). We co-cultured CAFs with bEND3-dnMAML cells or control bEND3 cells, and sorted CAFs for QPCR analyses. CAFs significantly upregulated CXCL10 expression upon co-culture with bEND3-dnMAML BECs compared to control bEND3 (Figure 4E). Culturing CAFs with conditioned media from bEND3-dnMAML cells was sufficient to induce fibroblast expression of *CXCL10* as well (Figure 4F), suggesting that loss of Notch signaling in ECs stimulates the production of soluble factor(s) that induce fibroblast *CXCL10*.

### Abrogation of Notch signaling in ECs reprogramed cancer-associated fibroblasts

To better understand how Notch signaling in tumor endothelium sculpts the local microenvironment, we further mined the scRNAseq results, and performed pathway analyses to assess how various tumor stromal cell types responded to the abrogation of Rbpj in ECs. Interestingly, of the cell types covered by scRNAseq, CAFs had the largest number of genes differentially expressed in Rbpj^iECKO^ versus control mice (Figure 5C), even exceeding the number of differentially expressed genes (DEGs) in ECs themselves.

In response to Notch deficiency in ECs, CAFs downregulated pathways involved in cell cycle and DNA replication (Figure 5E). Differential gene expression analyses of CAFs revealed a robust downregulation of molecules involved in cell cycle checkpoints, including *Gmnn, Ccnb2, Ccnd1, Cdkn2d, Cdk17, Cdca2, Cdca8, Cdc42ep3)* (Figure 5F). A number of molecules involved in DNA replication and cell division, including *Pold4, Top2a, Cenpa, Cenpe, Cenpf, Cenpm, Incenp, H2afz* were also downregulated, suggesting that CAFs halted proliferation in response to the loss of endothelial Notch signaling. To validate the transcriptomic analyses, we injected tumor-bearing Rbpj^iECKO^ and control mice with 5-bromo-2’deoxyuridine (BrdU). Sixteen hours later, approximately 6% of CAFs from control mice had incorporated BrdU. However, less than 2% of CAFs from Rbpj^iECKO^ mice were BrdU positive (Figure 5I), confirming that CAFs decreased proliferation in response to Notch deficiency in ECs.

Interestingly, CAFs also downregulated a number of molecules involved in smooth muscle contraction and Rho-mediated motility (Figure 5E and 5G), including *Acta2, Tpm4, Myl9, Cfl1, Pfn1*, *Tpm6, Mylk, Myl6, Vcl, Lmod1,* and *Tln,* indicating a loss of myofibroblast phenotype and function. Molecules associated with phenotypes of myofibroblasts, including *Ccn2 (Ctgf), Postn, Cxcl14, Serpine2, Timp1, Timp3, Mmp9, Col7a1, Col8a1, Col11a1, Col12a1, Tagln, Sdc1, Inhba* also decreased (Figure 5G). In addition, pathway analyses revealed a significant downregulation of TGFβ signaling, which is known to promote the myofibroblastic CAF phenotype (Biffi et al., 2019) (Figure 5E).

On the other hand, CAFs from Rbpj^iECKO^ mice had increased chemokine signaling, interferon signaling, and inflammatory responses (Figure 5D). Other than *Cxcl10*, we also saw increased expression of *Ccl7, Ccl8, Cxcl1, Cxcl2, Cxcl12, Il6, Jak1, Stat2, and Ackr3* (Figure 5H). In addition, molecules indicative of inflammatory CAF (iCAF) phenotypes, including *Dpp4, Ly6C, Cd248, Sfrp2, Scara3, and Has1* (Dominguez et al., 2020), were significantly upregulated in Rbpj-deficient mice (Figure 5H). Consistent with the transcriptomic data, CAFs from Rbpj^iECKO^ mice significantly upregulated the expression of inflammatory CAF marker Ly6C on the protein level (Figure 5J). Interestingly, one of the top pathways upregulated in CAFs was interleukin-1 (IL1) regulation of extracellular matrix (Figure 5D). Among the various tumor and stromal cell types, CAFs had the highest expression of Il1r1, the prototypical receptor that mediates IL1 signaling (data not shown). We also saw increased expression of *Il1r1* in CAFs of Rbpj^iECKO^ mice (Figure 5H). These observations were consistent with previous reports that IL1 antagonizes TGFβ to promote inflammatory phenotypes and functions in CAFs (Biffi et al., 2019). Collectively these data suggest that abrogation of endothelial Notch reprogrammed the tumor fibroblast compartment from myofibroblastic CAFs into pro-inflammatory CAFs to better engage anti-tumor immune responses (Elyada et al., 2019).

### Abrogation of Notch signaling in ECs unleashed interferon gamma-mediated induction of CXCL10

scRNAseq revealed a robust Rbpj^ECKO^-dependent upregulation of genes associated with type II interferon signaling in a wide spectrum of stromal cells, including not only CAFs, but also BECs, pericytes, TAMs, and granulocytes (Figures 6A–6D). Consistent with scRNAseq data, the amount of interferon gamma (IFNγ) transcript in the tumor tissue significantly increased in Rbpj^iECKO^ mice (Figure 6E). Because IFNγ is a well-documented inducer of CXCL10, we hypothesized that IFNγ mediated the CXCL10 induction and tumor inhibition in Rbpj^iECKO^ mice.

**Figure 6.**
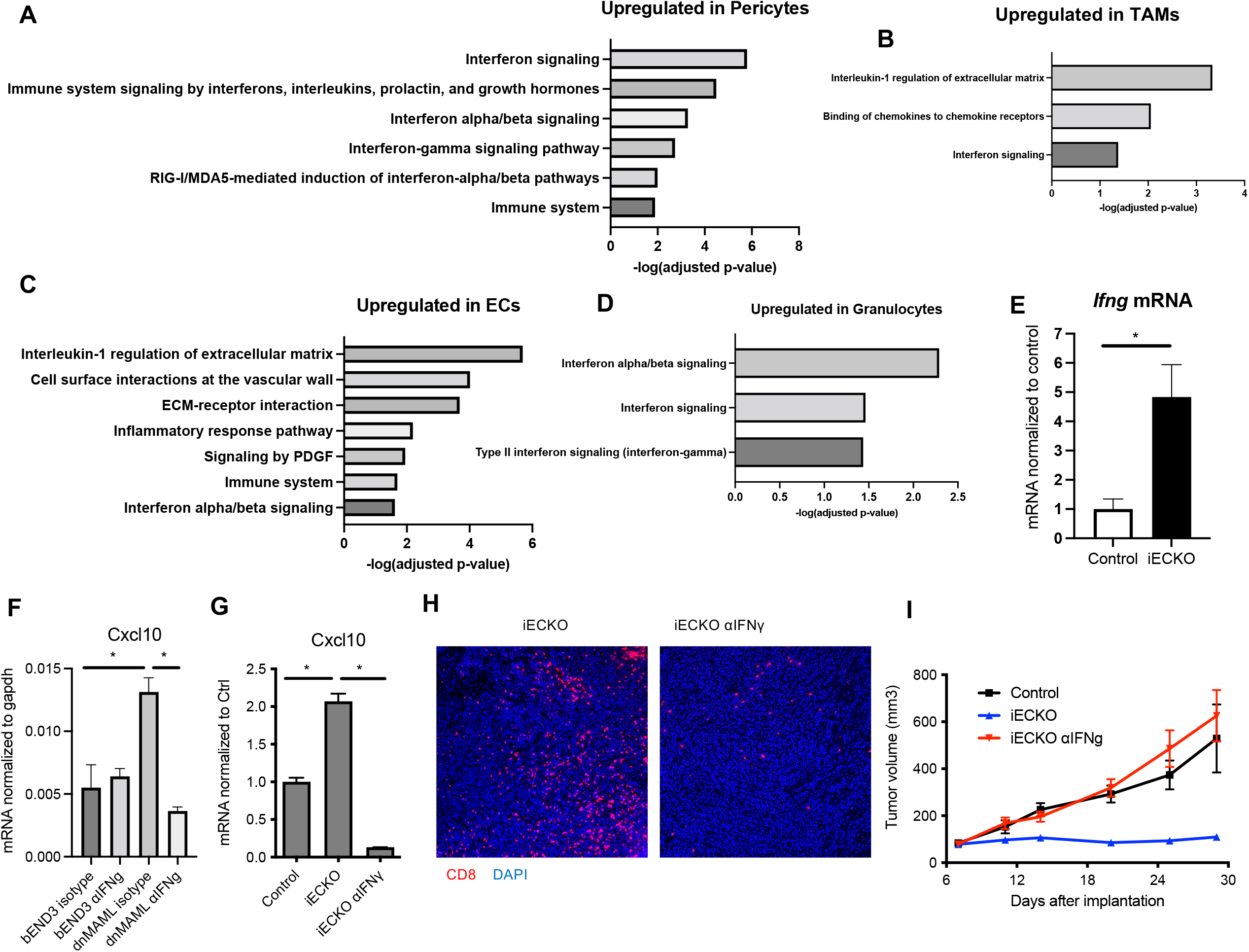
Abrogation of endothelial Notch unleashes IFNγ-mediated upregulation of CXCL10. A-D. Select BioPlanet pathways analyses of indicated tumor stromal populations that are upregulated by Rbpj^iECKO^. E. QPCR quantification of Ifng transcript in whole KP1 tissue lysates of control and Rbpj^iECKO^ mice. F. QPCR quantification of Cxcl10 transcript in CAFs treated with conditioned media from bEND3 or bEND3-dnMAML +/− αIFNγ. G. QPCR quantification of Cxcl10 transcript in whole KP1 tissue lysates of control and Rbpj^iECKO^ mice treated with αIFNγ. H. Immunofluorescence imaging of CD8+ cells in KP1 tumors of Rbpj^iECKO^ mice treated with αIFNγ. I. Tumor burden in (H).

We took advantage of our *ex vivo* model system to ask whether IFNγ was necessary or sufficient for the enhanced CXCL10 production by CAFs. As expected, recombinant IFNγ stimulated *CXCL10* transcription in CAFs, mimicking the effects of conditioned media from dnMAML-expressing Notch deficient ECs. Addition of anti-IFNγ neutralizing antibody suppressed the CXCL10 induction by conditioned media of Notch-deficient ECs (Figure 6F). These data suggest that Notch deficiency in ECs leads to increased CXCL10 production in an IFNγ-dependent manner.

Consistent with this model, *in vivo* administration of IFNγ neutralizing antibody dramatically reduced *Cxcl10* expression in tumor-bearing Rbpj^iECKO^ mice (Figure 6G). Suppression of CXCL10 correlated with reduced recruitment of CD8+ T cells in the tumor (Figure 6H). Importantly, neutralizing IFNγ reversed the suppression of tumor growth in Rbpj^iECKO^ PDAC-bearing mice (Figure 6I). Taken together, these data indicate that Notch signaling in tumor ECs inhibits the local production of CXCL10 by suppressing the IFNγ in the tumor environment. Abrogation of endothelial Notch signaling unleashed IFNγ responses to augment CXCL10/CXCR3-mediated T cell recruitment to inhibit tumor growth.

## Discussion

In this study we uncover a mechanism by which blood endothelial cells shield tumor cells from immune surveillance. We demonstrate that blood endothelial cells in PDAC upregulate canonical Notch signaling to inhibit the production of T cell recruitment chemokine CXCL10 by cancer-associated fibroblasts, thus protecting tumors from T cell-mediated attacks. Abrogation of Notch signaling in ECs reprogrammed myofibroblastic CAFs into inflammatory CAFs, unleashed Type II interferon responses in the tumor microenvironment, enhanced CXCL10 production by CAFs, and promoted the recruitment of anti-tumor T cells into the tumor tissue.

The abundance of anti-tumor T cells that infiltrate the tumor tissue closely correlates with cancer prognosis. The scarcity of tumor-infiltrating T cells for many patients, especially in those with certain types of cancers such as pancreatic ductal adenocarcinoma, remains a major challenge for cancer treatment. However, we only poorly understand the molecular and cellular events that sculpt an immunologically cold environment to protect tumor cells from immune surveillance. Accumulating evidence suggests that regulatory events within the immune system itself contribute to T cell scarcity. Innate immune cells, including tumor-associated macrophages (TAMs) and monocytic/granulocyte MDSCs, are well known for their ability to inhibit T cell responses through various mechanisms (Zhu et al., 2014). Insufficient dendritic cell presence in the tumor also causes defective presentation of tumor antigens and impairs activation of adaptive immunity that could inhibit tumor growth (Hegde et al., 2020).

How blood vessels promote the scarcity of tumor-infiltrating T cells is not well understood. Previous work showed that blood endothelial cells express Fas ligand on the cell surface to induce apoptosis in incoming T cells, thus preventing their function as tumor killers or suppressors (Motz et al., 2014). High levels of Fas ligand expression on endothelial cells correlate with low numbers of CD8+ T cells in multiple tumors, including bladder, renal, colon, and prostate cancers. Some studies also implicated the compromised functions of blood endothelial cells in mediating T cell adhesion to tumor vessels. In ovarian cancer models, for example, endothelin B receptor (ET_B_R) expressed on endothelial cells inhibits the level of intercellular adhesion molecule-1 (ICAM-1) on their cell surface, leading to decreased tumor-infiltrating lymphocytes (Buckanovich et al., 2008). Here we show that tumor endothelial cells also exert dramatic effects on stromal components of the tumor tissues to inhibit T cell trafficking, in a process dependent upon EC-intrinsic canonical Notch signaling. BECs in the pancreas enhance the activation of Notch signaling, making these cells interact with cancer-associated fibroblasts to modulate chemokine production in CAFs. The ability of CAFs to produce CXCL10, a chemokine essential for the recruitment of anti-tumor T cells, was inhibited by tumor endothelial cells. Abrogation of Notch signaling alleviated this suppression and unleashed the production of CXCL10 to attract T cell recruitment.

The chemokine receptor CXCR3, along with its ligands CXCL9/10/11, are well established members of the chemokine/receptor family that mediate Th1 CD4+ and CD8+ T cell trafficking (Groom and Luster, 2011a; Karin, 2018). This signaling axis has been shown to correlate with increased abundance of tumor-infiltrating T cells and better patient outcome in several types of cancers. In human PDAC patients, higher plasma concentrations of CXCL9 and CXCL10 in patients also correlate with improved overall survival (Qian et al., 2019). Abundance of CXCR3-expressing T cell infiltration has been shown to correlate with better responses to FOLFIRINOX (Peng et al., 2021). In mouse colorectal models, CXCR3 signaling was also shown to be required for responsivity to PD1 blockade (Chow et al., 2019). On the other hand, some human PDAC cells were shown to express CXCR3 that enhances tumor cell invasiveness (Hirth et al., 2020). However, in the mouse model that we used for this study, cancer cells and other stromal cells do not detectably express CXCR3, thus explaining its lack of tumor-promoting role. In these models and with intact Notch signaling in ECs, CXCR3 blockade did not dramatically change tumor size, suggesting that tumors suppressed the T cell recruitment function of CXCR3. However, the power of CXCL10-CXCR3 to recruit anti-tumor T cells was reactivated upon the abrogation of endothelial Notch signaling. It remains to be seen whether such alleviation could be harnessed in human PDAC to augment anti-tumor immunity and benefit patient, without engaging the confounding pro-tumor effects on carcinoma cells.

Short-term homing assay suggests that engagement of CXCR3 signaling can recruit circulating T cells that originate from draining lymph nodes, thus converting “cold” tumors into “hot” tumors. It is important to note that CXCR3 could also orchestrate intratumor localization of T cells pre-existing in the tumor tissue, thus subverting the “immune excluded” environment to augment anti-tumor immune responses (Chow et al., 2019). Along the same line, in subsets of human colorectal cancers, scRNAseq revealed foci of activated IFNγ + T cells surrounded by myeloid cells expressing CXCR3 ligands (Pelka et al., 2021), suggesting a role of this chemokine receptor in organizing the spatial localization and functions of T cells in addition to their recruitment.

Interferon gamma acts as an important bridge between the abrogation of Notch signaling in ECs and the expression of CXCL10 in CAFs. Neutralization of IFNγ not only reversed the increase in CXCL10 seen in the tumors of Rbpj^iECKO^ mice, but also abolished the anti-tumor effect brought forth by endothelial Notch deficiency. However, ECs do not appear to be prominent sources of IFNγ in the tumor microenvironment; scRNAseq suggest that T cells and NK cells are the most potent IFNγ producers. Molecular mechanisms by which endothelial Notch inhibits IFNγ also need to be determined. A better analyses of the EC secretome is also needed to understand how Notch directs EC interaction with the tumor stroma to shape the local cytokine/chemokine milieu and regulate leukocyte recruitment.

Cancer-associated fibroblasts are well known for modulating the access of T cells to the tumor (Feig et al., 2013; Sahai et al., 2020). CAFs have been demonstrated to shape an “immune excluded” environment, where T cells are present in the organ but blocked from accessing malignant cells. The underlying mechanisms include the production and secretion of extracellular matrix by CAFs, creating a fibrotic barrier that could physically shield T cells. CAFs are also notorious for producing myeloid cell-recruiting cytokines and chemokines that enhance the abundance of TAMs, MDSCs, and regulatory T cells to render T cells dysfunctional (Feig et al., 2013; Kumar et al., 2017; Takahashi et al., 2017; Tan et al., 2011; Zhao et al., 2020). Various cell types within the tumor microenvironment, such as tumor cells and myeloid cells, shape the functions of CAFs, and CAFs in turn crosstalk with these cells to regulate tumor growth. Our data show that tumor blood vessels also shape CAF functions. This mechanism differs from those by which CAFs modulate an excluded environment. Instead of creating fibrotic shields, CAFs modulate their chemokine profile to alter T cell homing from draining lymph nodes (Elyada et al., 2019); this type of modulation depends on soluble factors that include IFNγ. In addition, Notch-deficient ECs secrete other cytokines that not only synergize with IFNγ to induce CXCL10 (data not shown). Other modes of crosstalk await elucidation, providing a potentially fertile ground for future studies on EC-CAF-leukocyte interactions.

## Acknowledgements

The authors acknowledge support from NIH R01 grants CA228019 and AI130471 (E.C.B.), Cancer Research Institute Irvington Postdoctoral Fellowship (Y.Z.), and Tobacco-Related Disease Research Program Postdoctoral Fellowship (M.X.). Sequencing was performed at Stanford Functional Genomics Facility with instrumentation funded by NIH S10OD025212 and 1S10OD021763. The KP1 and KP2 cell lines were kind gifts from David G. DeNardo. The authors would like to thank all members of the Butcher Laboratory for input on this work.

## STAR Methods

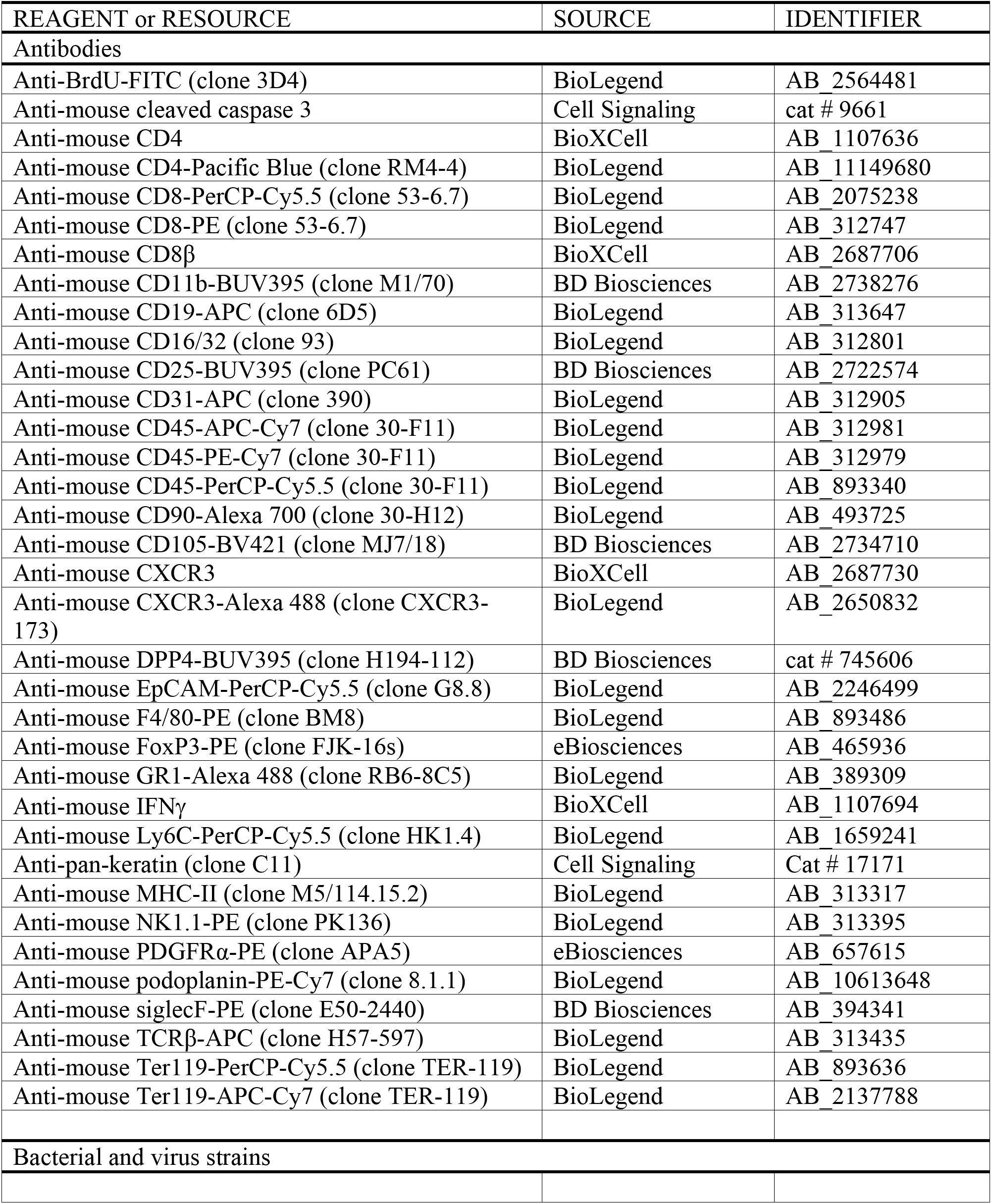
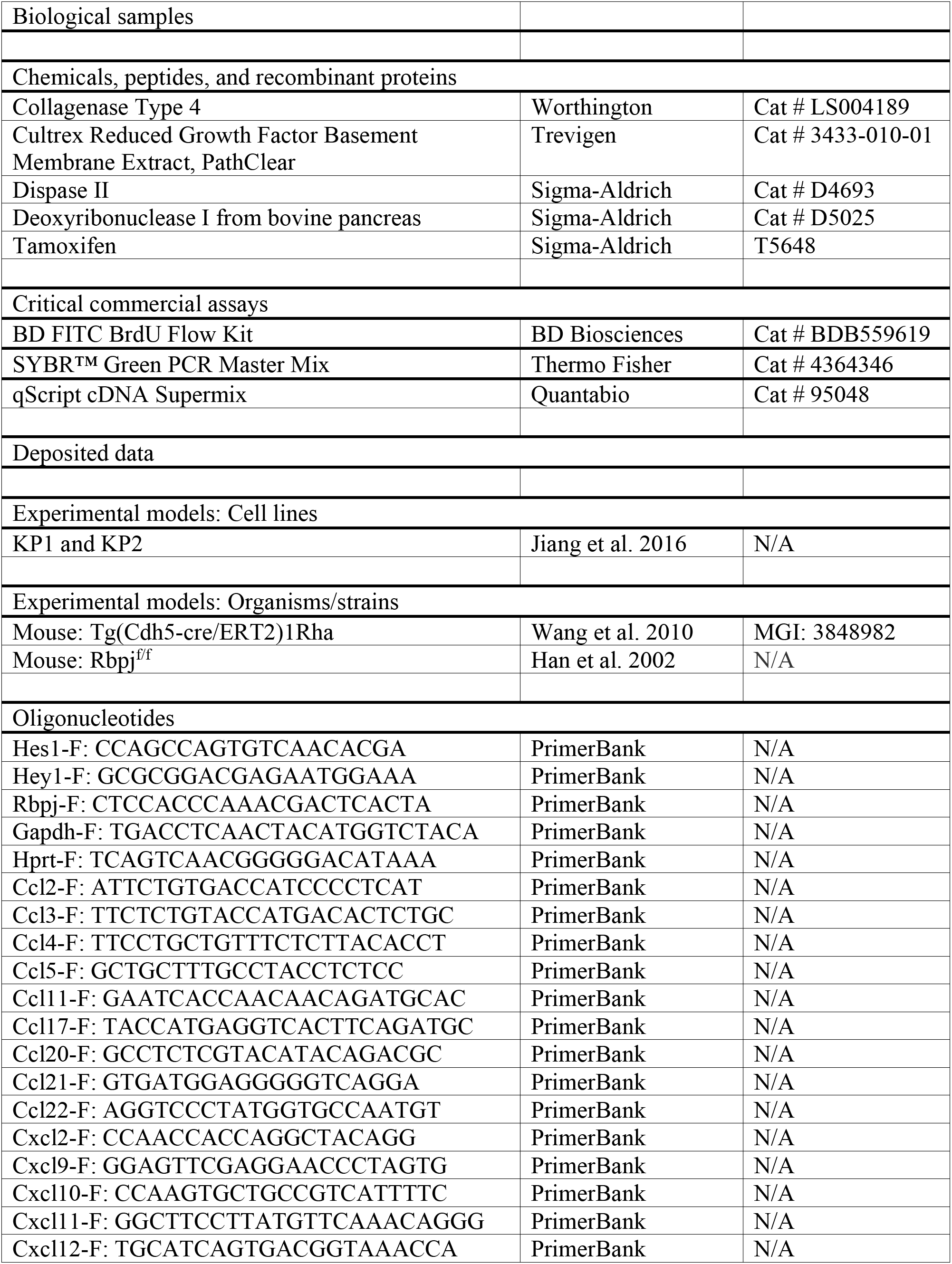
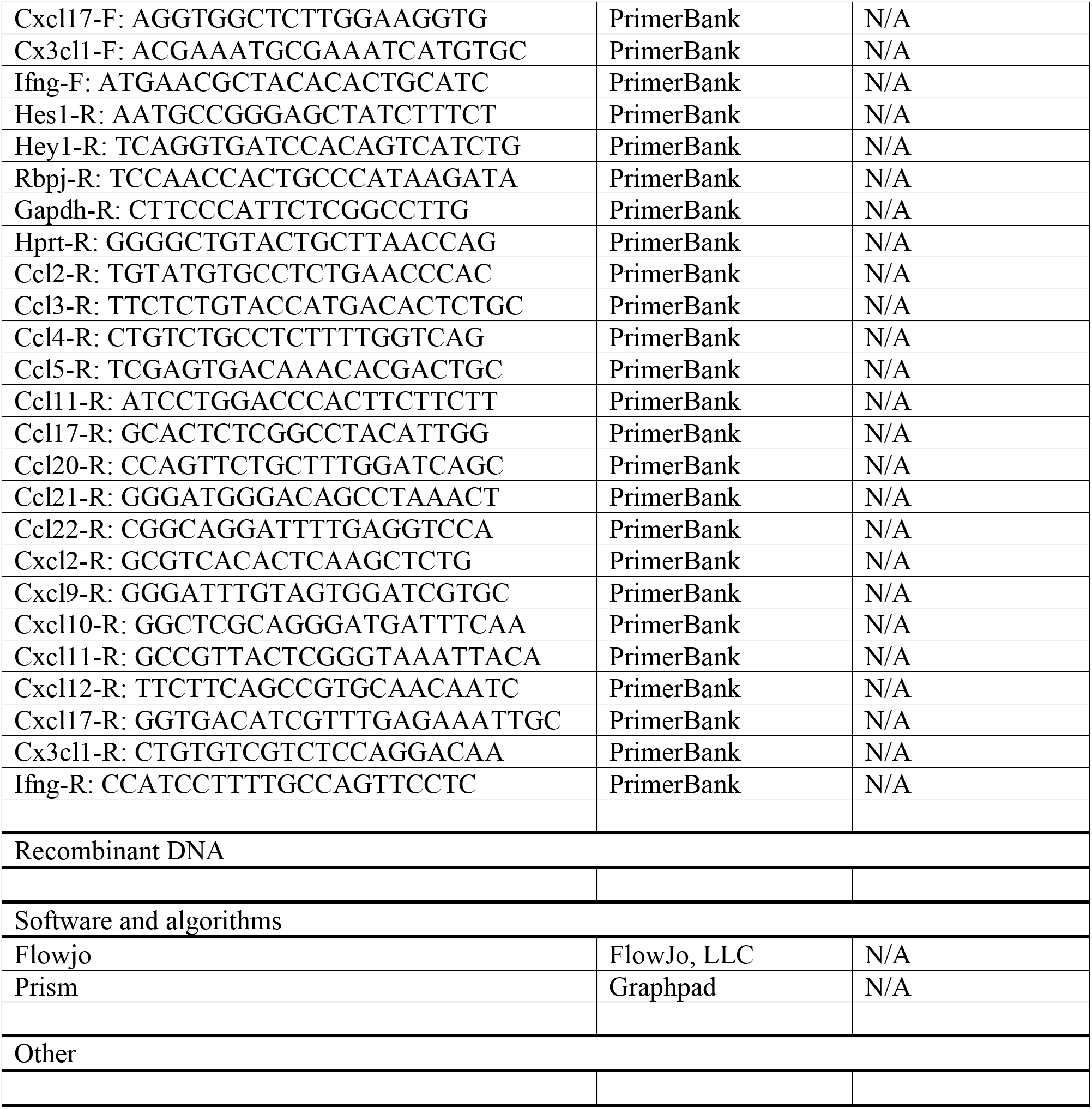
Key Resources Table

### Lead Contact

Further information and requests for resources and reagents should be directed to and will be fulfilled by the lead contact, Eugene C. Butcher.

### PDAC Model and Treatment

To establish KPC models, between 20,000 and 100,000 KP1 or KP2 cells in 50 μL of Cultrex (Trevigen) were injected orthotopically into the pancreas or subcutaneously into the left flank of 2 to 6-month-old mice. Once tumors became palpable orthotopically or reach 0.5cm in length subcutaneously, Cdh5-Cre^ERT2^; Rbpj^loxP/loxP^ and control mice (Cdh5-Cre^ERT2^ only or Rbpj^loxP/loxP^ only) were treated with two doses of 150ug/kg tamoxifen. Concurrent with the first tamoxifen injection, we also started treating mice intraperitoneally (i.p.) with antibodies (BioXCell) against CXCR3 (CXCR3-173, 10mg/kg), IFNγ (XMG1.2, 200mg), CD4 (GK1.5, 200mg), CD8β (53-5.8, 200mg), or isotype controls, including Armenian hamster IgG and rat IgG1 (HRPN) or IgG2b (LTF-2), twice a week. Tumor volume was quantified as ½ x length x width^2^. Survival was scored when mice lost 15% of body weight, tumors reached 1.5cm in diameter, or upon death. All animal work was approved by Institutional Animal Care and Use Committee at the Veterans Affairs Palo Alto Health Care System.

### Tissue Isolation and Flow Cytometry

Tumor tissues were manually minced and digested in 20 mL of Hanks Balanced Salt Solution (HBSS) (Thermo Fisher) containing 500U/mL collagenase D (Worthington Biochem), 20 μg/mL DNase I (Sigma), and 2% fetal bovine serum (FBS) for 30 min at 37C with constant stirring. Normal pancreas was digested for 15 min. Digestion was quenched by ethylenediaminetetraacetic acid and filtered through 40 μm Nylon mesh, pelleted through centrifugation (750g for 5 min at 4C), and resuspended in phosphate buffered saline (PBS).

Single cell suspensions were incubated in PBS with anti-mouse CD16/CD32 antibodies (1/200) (eBioscience) and Zombie NIR™ Fixable Viability dye (1/500) (BioLegend) for 10 min, pelleted by centrifugation, and subsequently labeled with 100-200 μL of fluorophore-conjugated anti-mouse antibodies at pre-determined dilutions for 20 min on ice, and washed with staining buffer (PBS with 1% FBS). For proliferation assays mice were injected with BrdU (4 mg) i.p. 16 hr prior to sacrifice. BD Bioscience Cytofix/cytoperm kit was used to stain for BrdU. Data were acquired on LSR-Fortessa (BD Biosciences) and analyzed using FlowJo software.

### Lymphocyte Homing Assay

We implanted 50,000 KP1 tumor cells orthotopically in the pancreas of donor (Cre^Neg^, Rosa26-mTmG) and recipient (Cdh5-Cre^ERT2^; Rbpj^loxP/loxP^ and control) mice. When tumors reached approximately 1cm in diameter, we injected recipient mice with tamoxifen (4mg/dose) on two consecutive days. Twenty-four hours later, we isolated the hepatic, portal, and mesenteric lymph nodes of donor mice, crushed through 40 filters, treated with red cell lysis buffer (BioLegend), washed and resuspended in 200 μL of PBS, and injected intravenously into the recipient mice. Approximately 10^7^ live cells were injected into each recipient. Donor cells were confirmed to be tdTomato+. Sixteen hours after injection, we sacrificed recipient mice for flow cytometry analyses.

### RNA Isolation and Quantitative PCR

RNA was isolated using E.Z.N.A. Total RNA kit (Promega). cDNA was synthesized using qScript cDNA SuperMix (Quantabio) following manufacturer instructions. Quantitative PCR was performed using SYBR Green Master mix (Thermo Fisher). Primers were either designed using Primer3 or referenced from PrimerBank and synthesized by Integrated DNA Technologies.

### Immunofluorescence Imaging

Fresh tumor tissues were fixed in 2% formaldehyde overnight, incubated with 30% sucrose overnight, and embedded in OCT compound (Sakura Finetek). Six μm cryosections were blocked with PBS and 5% serum (1hr at room temperature), incubated with rabbit anti-cleaved caspase 3 (Cell Signaling) overnight, and then with Alexa488-conjugated goat anti-rabbit IgG (ThermoFisher), PE-CD8 (BioLegend), and Alexa647-conjugated anti-pan-keratin (C11, Cell Signaling) antibodies for 1hr at room temperature. Tissues were washed with PBS containing 0.05% Tween-20 between incubation steps, and mounted in DAPI-containing Fluoromount-G (SouthernBiotech) for imaging on Apotome (Zeiss).

### Single-cell RNA sequencing

KP1 tumor cells (100,000) were established in Rbpj^iECKO^ and control mice (8-9 mice/group). Mice were treated with 150mg/kg of tamoxifen on Day 10 and 13, and sacrificed on Day 14. Cells from Rbpj^iECKO^ and control mice were isolated in parallel and stained with TotalSeq-A hashtag antibodies (BioLegend) simultaneously with flow antibodies. Populations of interest were then sorted and processed for scRNAseq using Chromium Single Cell 3’ Library and Gel Bead Kit v3.1 (10x Genomics) according to manufacturer guidelines. Male and female mice from Rbpj^iECKO^ or control cohort were processed together and resolved post-sequencing. Libraries were sequenced on NovaSeq 6000 (Illumina) at Stanford Functional Genomics Facility. Cell Ranger (v6.0.1, 10x Genomics) was used to align reads to the mm10 reference genome and perform quality control. Count data were processed with the Seurat package (v4.0.2). Raw counts were log normalized and 2000 most variable genes were identified based on a variance stabilizing transformation. Principal Component Analysis (PCA) dimensionality reduction was performed using the variable gene sets. Cell clusters were determined using a Shared Nearest Neighbor (SNN) modularity optimization-based clustering algorithm of the Seurat “FindClusters” function, and were visualized with Uniform Manifold Approximation and Projection (UMAP). Contaminating cells were removed manually. To recover gene-gene relationships that are lost due to dropouts, missing gene expression data from log normalized count data was imputed using the MAGIC (Markov Affinity-based Graph Imputation of Cells) algorithm with optimized parameters (*t* = 2, *k* = 9, *ka* = 3) (van Dijk et al., 2018) (ref). Imputed data were used for visualization of single-cell gene expression in dot plots.

Differential gene expression analysis was performed using DESeq2 (v1.30.1) on subset mean gene expression values (Love et al., 2014). Gene enrichment analysis of DEGs with BioPlanet databases was performed using Enrichr (Kuleshov et al., 2016).

