## Supplemental Figure for "Notch signaling in tumor vasculature programs cancer-associated fibroblasts to suppress anti-tumor immunity"

**Figure S1. Endothelial Notch signaling promotes tumor growth in murine tumor models**

Weights of subcutaneous KP1 or KP2 tumors in control and $Rbpj^{iECKO}$ mice.

**Figure S2. Gating strategy for immune cell subsets**

Representative flow cytometry plots for identification of lymphocytes, macrophages, eosinophils, monocytes or monocytic myeloid-derived suppressor cells (MDSCs), and granulocytes or granulocytic MDSCs.

**Figure S3. Endothelial Notch signaling inhibits the infiltration of anti-tumor T cell in multiple tumor models**

A. Flow cytometric quantification of indicated lymphocytes in subcutaneous KP1 tumors in control and $Rbpj^{iECKO}$ mice one day after tamoxifen induction.

B. Flow cytometric quantification of myeloid cells (A) and T cells (B) in orthotopic KP2 tumors of control and $Rbpj^{iECKO}$ mice.

C. Representative flow cytometry plots and quantification of T cells in orthotopic KP1 tumors from control and $Rbpj^{iECKO}$ mice treated with CD4/CD8 depleting antibodies.

D. Tumor burden of mice in (C).

**Figure S4. CXCR3 is expressed on T cells, but not on tumor cells and other stromal cells**

Representative histogram of CXCR3 staining in indicated cell populations by flow cytometry. Isotype controls were shown as shaded gray.

**Figure S5. Sorting strategy for BECs, CAFs, and tumor cells.**

Representative flow cytometric plots for identification of blood endothelial cells (BECs), cancer-associated fibroblasts (CAFs), and tumor cells.

Supplemental Figure 1. Endothelial Notch signaling promotes tumor growth in murine tumor models

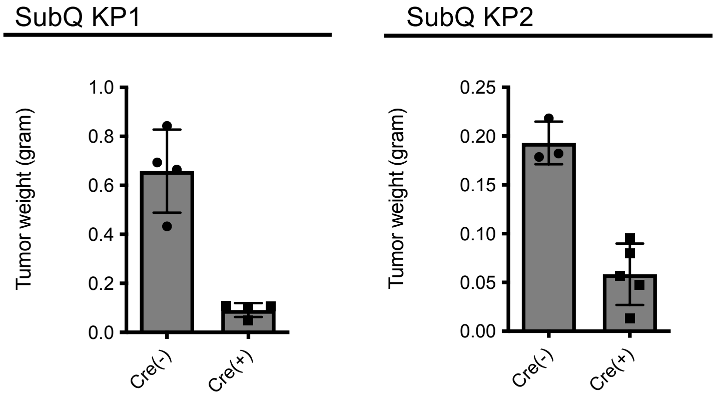

Supplemental Figure 2. Gating strategy for immune cell subsets

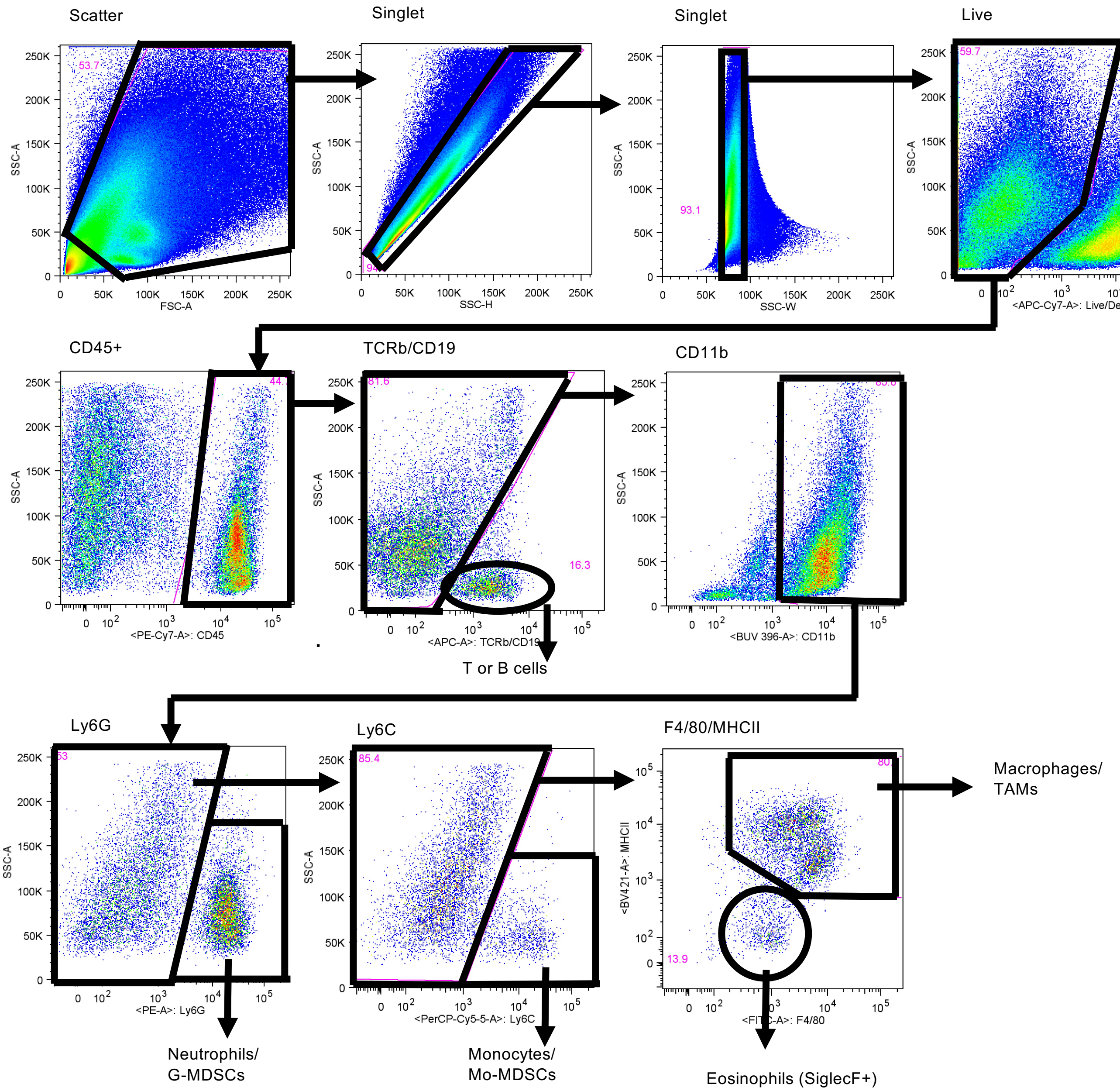

Supplemental Figure 3: Endothelial Notch signaling inhibits the infiltration of anti-tumor T cell in multiple tumor models

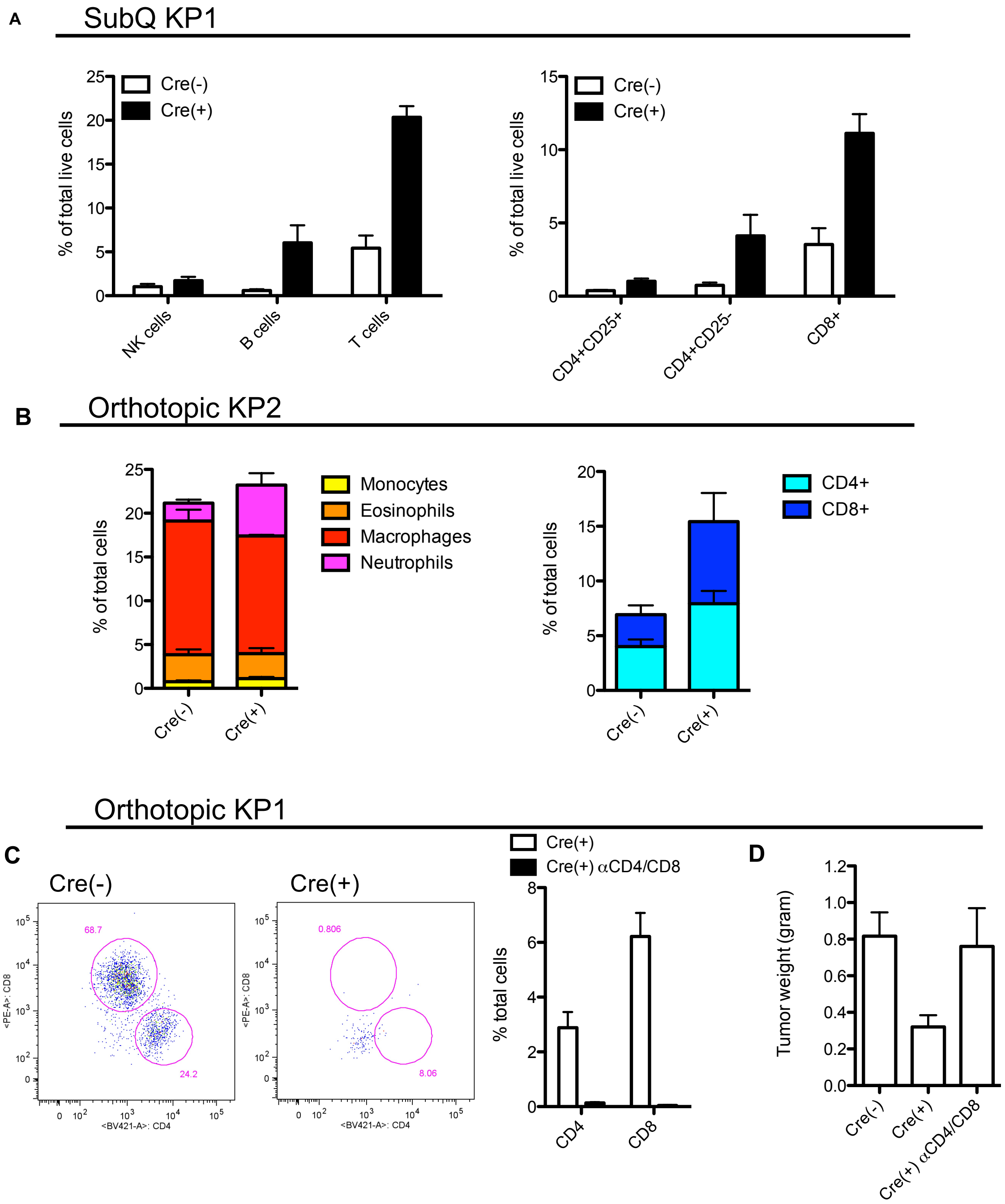

Supplemental Figure 4. CXCR3 is expressed on T cells, but not on tumor cells and other stromal cells

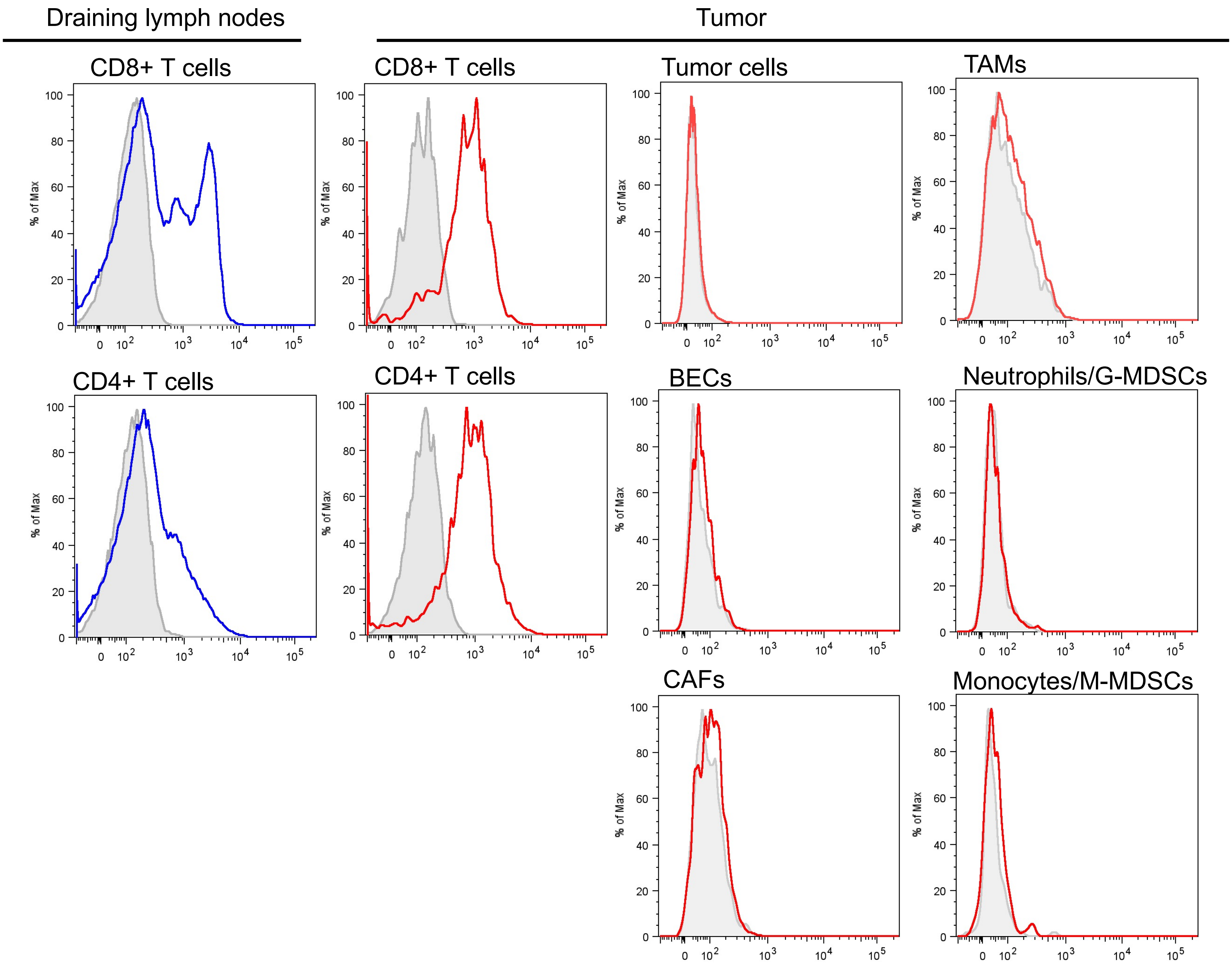

Supplemental Figure 5. Sorting strategy for BECs, CAFs, and tumor cells

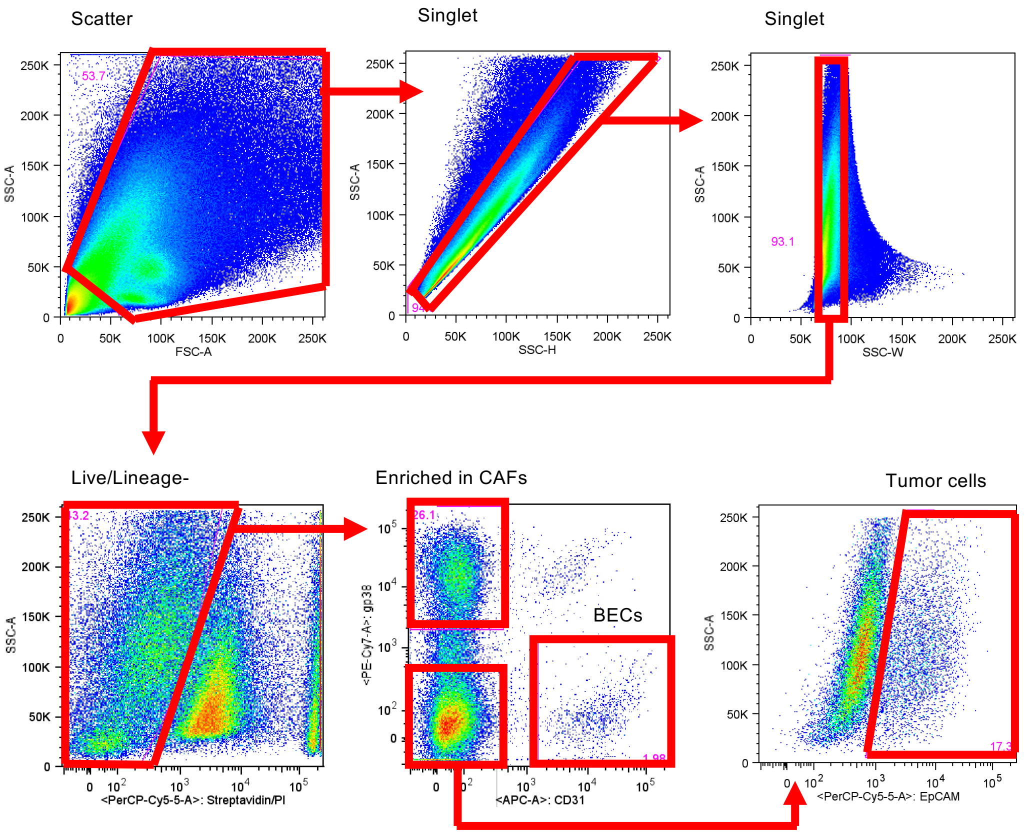
